# Differences in Methanotrophic Community Structure in Two Methane-Rich Habitats: Oil Natural Gas Field & Paddy Field

**DOI:** 10.1101/2024.06.03.597264

**Authors:** Akanksha Verma, S.S. Maitra

## Abstract

Methanotrophic bacterial isolates were identified in this study using the molecular detection method, isolated using microbiological techniques, and studied their cellular shape using atomic force microscopy. Two methanotrophic bacterial species belonging to the *Methylocaldum* and *Methylomonas* genera were provisionally designated as Isolate 1 and Isolate 5, thus isolated from the Oil-Natural Gas Field and Paddy Field, respectively. The Oil-Natural Gas Field Isolate 1 showed 91.82-97.25% sequence homology to the reference Methanotrophic species, whereas Paddy Field Isolate 5 showed 79.72-84.99% sequence homology to the reference *Methylomonas* species in the NCBI database. As per the phylogenetic analysis, Oil-Natural Gas Field Isolate 1 and Paddy Field Isolate 5 are possibly new species of *Methylocaldum* and *Methylomonas* genus, respectively. In addition, the microscopic study also supported the molecular identification and phylogenetic analysis of isolated species by showing the cocci and rod shapes for the Oil-Natural Gas Field Isolate 1 and Paddy Field Isolate 5, respectively.

## 1. Introduction

Methanotrophs, a methane-oxidising bacteria, are a distinct group of methanotrophic bacteria that rely solely on methane (CH_4_) for carbon and energy sources (1). Aerobic methanotrophic bacteria are categorised within the phyla Proteobacteria and Verrucomicrobia. In contrast, the process of anaerobic methane oxidation is facilitated by newly identified anaerobic methanotrophs (2). They are present in both the bacterial and archaeal domains. (3) found almost everywhere in nature, and they have been isolated from a wide range of habitats, including soils, peatlands, rice paddies, sediments, freshwater and marine systems, acidic hot springs, mud pots, alkaline soda lakes, cold settings, and tissues of higher species (4). It plays a pivotal role in the methane cycle by maintaining the balance of atmospheric methane in the environment (5). Methanotrophs can use the metabolic by-products of methane oxidation (methanol, formaldehyde, and occasionally other C_1_ compounds) (6). Methanotrophic bacteria are classified into three groups: Type I, Type II, and Type X, where they differ in terms of their phylogenetic affiliation, such as *gammaproteobacterial* (Type I and X) versus *alphaproteobacterial* (Type II) and a variety of biochemical properties. A fragment of the *pmoA* gene, encoding the active-site subunit of particulate methane monooxygenase (pMMO), is commonly used to identify methanotrophic bacteria in soils via the cultivation-independent method. Except for *Methylocella palustris* and *Methylocella silvestris*, this marker gene is found in all known methanotrophic bacteria (7). The particulate methane monooxygenase (pMMO), soluble methane monooxygenase (sMMO), and methanol dehydrogenase (MDH) are highly conserved enzymes found in Methanotrophs. The enzyme pMMO is expressed in all methanotrophs, whereas sMMO is expressed only in Type I, II and *Methylococcus capsulatus* (8). Both these enzymes are involved in the conversion of methane into methanol. The enzyme MDH is expressed in all gram-negative methanotrophic bacteria (9), which converts methane into formaldehyde. Therefore, detecting these enzymatic activities or the genes that encode these enzymes may allow for detecting all known methanotrophs (Jhala et al., 2014). Type I methanotrophs utilise the Ribulose Monophosphate (RuMP) pathway, whereas Type II and Type X follow the Serine pathway (9).

CH_4_ is a potent greenhouse gas, and methanotrophs help to reduce the amount of CH_4_ released into the atmosphere (9). Methanotrophs are essential in regulating CH_4_ emissions from paddy fields (11). Only a portion of the CH_4_ produced in rice field soil is released into the atmosphere; the rest is oxidised by methanotrophic bacteria that live in oxic niches in flooded fields (i.e., the surface soil layer and the rhizosphere). When anoxic rice field soil slurries are aerated, they transition from CH_4_ production to CH_4_ consumption (Henckel et al., 1999). Therefore, the oil field and paddy field were the prime choices in this study for the isolation of methanotrophic bacteria.

Microbiologists have identified the presence of uncultured organisms *in situ* using molecular biological tools and the 16S rRNA gene as markers. These 16S rRNA-based methods have resulted in estimates of prokaryotic biodiversity and surveys of the microbial community structure of various environments (13). Here, two different Methanotrophs of the genus *Methylocaldum* and *Methylomonas* were identified using culture-independent molecular techniques such as PCR amplification and phylogenetic tree analysis. *Methylomonas* bacterial species were isolated from the Paddy Field (PF) soil, sampled from a small rural village, Bara, located in the Banka district of Bihar, India and *Methylocaldum* bacterial species were isolated from the Oil-Natural Gas Field (ONGF) soil sampled from Jorhat, Assam, India. Both the bacterial species were characterised by their physiological and morphological characteristics along with the phylogenetic analysis (14)(15)(16). Furthermore, the microscopic analysis of isolated colonies from both samples revealed the difference in their cellular shape by Atomic Force Microscopy (AFM). The identified bacteria differed in their membrane structure, carbon fixation pathways, and phylogeny relationships.

## 2. Materials & Methods

### 2.1. Study sites & and soil sampling

Soil samples were systematically collected at a depth of 8-10 cm from the surface of the designated collection site. The collection process involved obtaining soil samples from eight distinct locations originating from four separate areas affiliated with Oil & Natural Gas (Borholla GGS, Arkin basin drilling services, Nambar GGS I, and Lahoti MCD Dumping site) situated in Jorhat, Assam, India (26°45’25.5” N 94°14’41.2” E) (refer to **Table 1**). Official permissions were duly acquired from the Oil & Natural Gas Corporation (ONGC) and the Institute of Biotechnology & Geotectonic Studies (INBIGS) in order to conduct the collection of soil samples from the oil fields. Furthermore, two paddy fields were specifically chosen for soil sample collection, namely, (1) Chhap, Muzaffarpur, Bihar, and (2) Bara, a small village located in the Banka district of Bihar, India (24°57’46.4868” N 86°37’34.4238” E) (refer to **Table 1**).

**Table 1:**
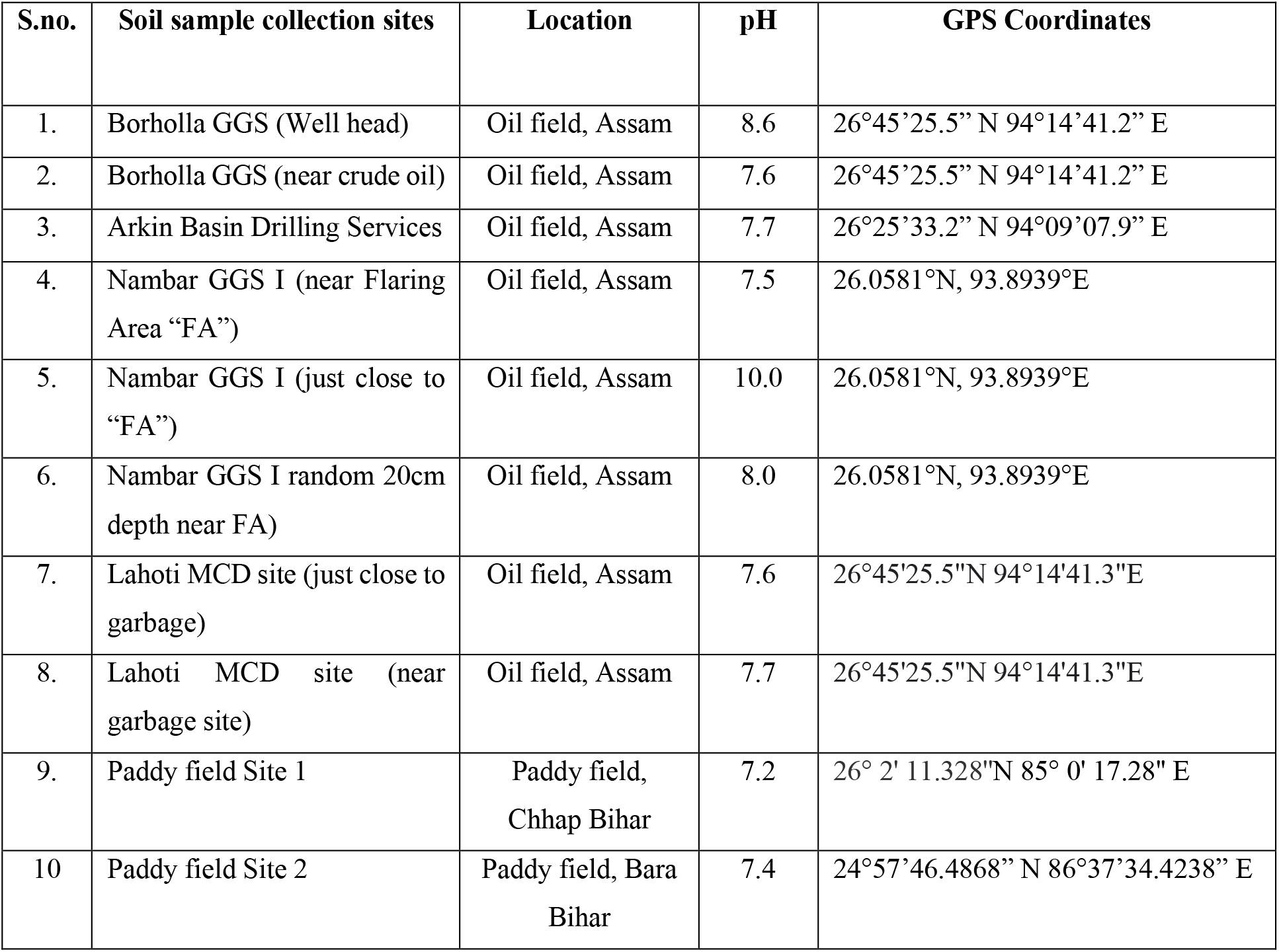
ONGF and PF soil sample collection sites and their GPS coordinates.

Immediately after collection, all soil samples were promptly transported to the laboratory within a maximum time span of six hours. Upon arrival, the samples were immediately processed, and the remaining were stored at a temperature of −20°C. Subsequently, 100 grams of each soil sample were transferred to sterile blue-capped bottles containing 100 ml of distilled water. These bottles were subjected to shaking for a duration of 15 minutes on an orbital shaker. The contents of the bottles were then filtered using Whatman Filter paper one and subsequently stored at a temperature of 4°C. In order to facilitate further analyses, all soil samples were subjected to serial dilution up to a ratio of 10^−4^ and enriched in Nitrate Mineral Salt (NMS) medium (17) within serum bottles.

### 2.2. Medium and culture conditions for bacterial enrichment and isolation

NMS medium was prepared using MgSO_4_, KNO_3_, CaCl_2_, trace element solution, vitamin stock solution, and iron stock solution as described previously (18). The pH values of the soil samples obtained from the oil fields ranged from 7.4 to 7.6, 8 to 8.6, and 10, while the pH of the paddy field soil was 7.4. To maintain consistent conditions, sodium phosphate buffers at pH 7.4, 8.8, and 10 were prepared as suitable buffer solutions. After autoclaving, the sodium phosphate buffer at pH 7.4, vitamin stock solution, and CuCl_2_ were added to the NMS medium.

Following the filtration of soil samples using a Whatman filter paper 1 by dissolving the soil in distilled water, the filtrate was utilized as a starter culture for inoculating bacterial cultures. Autoclaved NMS media (10 ml) (17) were then transferred into serum bottles along with 1 ml of filtered soil sample. Methane was introduced into the serum bottle inoculated with the culture using a 10 ml syringe at a ratio of methane to air of 1:4 (the methane to air ratio is maintained by evacuating the air from the serum bottles by using an aseptic syringe). The serum bottles were covered with butyl rubber caps and sealed using aluminium crimp seals. The cultures were subsequently incubated at a temperature of 30°C and shaken at 150 rpm for a duration of five days (~120 hrs) (18). On the 5^th^ day, the cultures showed visible turbidity. Therefore, cultures were fully grown and ready for further processing.

The grown cultures were streaked onto NMS agar plates and placed in an anaerobic jar containing 20% methane. The plates were then incubated at a temperature of 30°C for a period of 10 to 15 days until bacterial colonies became visible. Methane was regularly added at intervals of 4 days until the appearance of bacterial colonies. Once growth was observed on the NMS plates, individual isolated colonies were patched (The method involves utilizing a sterile toothpick to lift an entire bacterial colony (or a portion thereof) and move it onto the surface of a new agar plate) onto another NMS agar plate and again placed in an anaerobic jar with 20% methane (20% methane was maintained through evacuating the vacuum from the anaerobic jar through a vacuum pump). This plate was incubated at 30°C until proper bacterial growth was achieved, which typically took around 10 days. Similarly, methane was added at regular intervals of 4 days until bacterial colonies became visible. Once growth was observed on the patched plates, a small inoculum from each colony was selected and enriched in NMS broth medium within serum bottles with a methane-to-air ratio of 1:4. The inoculated cultures were then incubated at 30°C and shaken at 150 rpm for a duration of five days (~120 hrs) to cultivate the methanotrophic bacteria.

### 2.3. Pure-Culture preparation

To isolate and obtain a single colony of methanotrophic bacteria, the mixed bacterial culture from both the oil field and paddy field was further grown in an NMS medium and streaked on NMS agar plates, followed by repeated culturing and streaking until single colonies were obtained. The isolated colonies from each sample were inoculated in NMS broth media for further studies, such as sequencing and microscopic studies for phylogenetic and morphological analysis, respectively.

### 2.4. DNA extraction

The genomic DNA was isolated from the bacterial culture enriched in an NMS medium using a Genomic DNA Isolation kit by GCC Sure GCC Biotech. The concentrations of extracted DNA samples were determined using a NanoDrop ND-2000 spectrophotometer (*Thermo Fisher Scientific, USA*) and quality by running it on 0.8% Agarose gel electrophoresis at 100V for 40 minutes). The extracted DNA samples ranged from 150-210 ng/μl.

### 2.5. Polymerase Chain Reaction

The genomic DNA extracted from both soil samples underwent subsequent processing for PCR amplification using universal primers (refer to **Table 2**) targeting the 16S rRNA gene to facilitate the identification of bacterial species, as well as primers specific to the particulate methane monooxygenase (pMMO) gene for the detection and characterisation of methanotrophic bacteria. For the amplification of the 16S rRNA gene, the PCR protocol involved an initial denaturation step at 95°C for 5 minutes, followed by 35 cycles of denaturation at 94°C for 30 seconds, annealing at 54°C for 45 seconds, extension at 72°C for 1 minute, and a final extension step at 72°C for 1 minute. The PCR reaction was then cooled down to 4°C. To identify and characterise the methanotrophic community within the soil samples, the pMMO gene was targeted. The PCR amplification protocol for the pMMO primers included an initial denaturation step at 94°C for 1 minute, followed by 30 cycles of denaturation at 94°C for 1 minute, annealing at 55°C for 1 minute, extension at 72°C for 1 minute, and a final extension step at 72°C for 5 minutes. The PCR reaction was then cooled down to 4°C. PCR products obtained from both amplifications were subsequently purified using a PCR purification kit (*Fermentas, UK*) to remove any unwanted residues and contaminants, ensuring high-quality DNA for subsequent sequencing analysis.

**Table 2:**
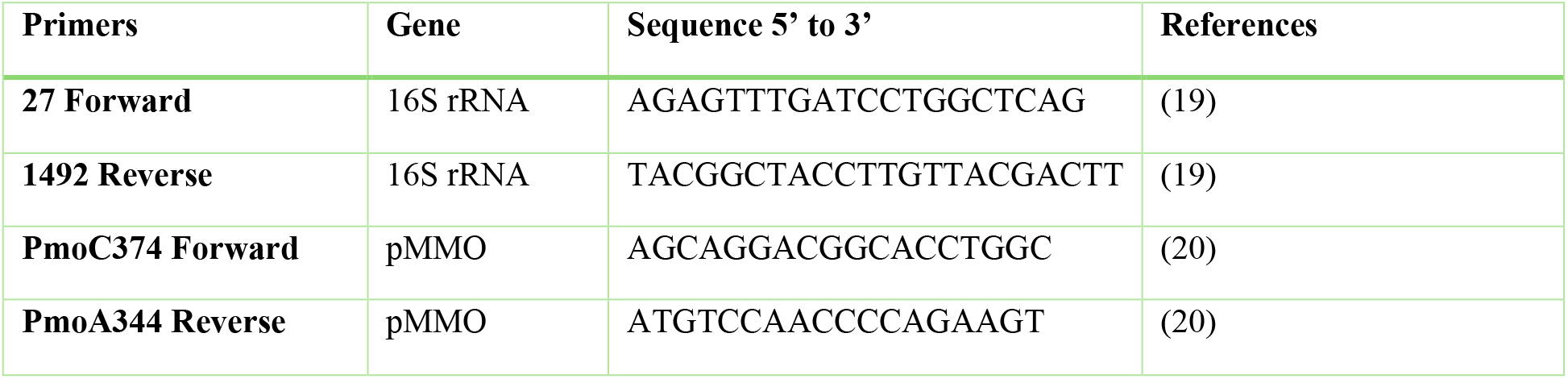
List of primers used in PCR amplification.

### 2.6. Phylogenetic analysis

The obtained partial 16S rDNA sequences from this study were subjected to a similarity search in the NCBI database using the BLASTn program (Basic Local Alignment Search Tool for nucleotide). The query sequence was copied and pasted into the BLASTn search tool, and the search was submitted. Among the 100 sequences obtained, those exhibiting a similarity above 90% were selected for further analysis. These sequences were aligned using the MUSCLE algorithm in MEGA X software version 11 (21).

To assess the phylogenetic relatedness among the obtained clones, a Maximum Likelihood tree was constructed. The UPGMA (Unweighted Pair Group Method with Arithmetic Mean) model was employed, and the tree was generated with a bootstrap value of 1000. During the analysis, any positions in the dataset containing gaps or missing data were excluded using the complete deletion option to ensure accurate and reliable results.

### 2.7. Atomic Force Microscopy (AFM)

Atomic Force Microscopy (AFM) is a powerful imaging technique that enables high-resolution examination of surfaces at the nanometer (nm) scale. It operates by mechanically probing the surface and can visualize cells and biomolecules without the need for chemical treatments. In order to ensure accurate imaging, it is essential to firmly immobilize the samples on the mounting surface to prevent displacement caused by the forces exerted by the scanning AFM cantilever tip. The immobilization process should be minimally invasive to avoid interfering with the metabolic processes or biochemical properties of the samples.

To capture images of negatively charged methanotrophic bacteria, freshly cleaved mica surfaces are coated with porcine (pig) gelatine as described by (21). A gelatine solution was prepared by dissolving 0.5 gm of gelatine in 100 ml of hot distilled water, thoroughly mixing, and then cooling down to a temperature of 60-70 °C. The gelatine solution was then transferred into a small beaker (20 ml), into which the cut and cleaved mica sheets were immersed. For mica coating, the mica sheets were dipped into the gelatine solution and allowed to dry on a paper towel. To prepare the bacterial sample, a bacterial pellet was obtained by centrifuging 1 ml of the culture at 4500 rcf for 5 minutes. The pellet was washed twice with deionized water and then resuspended in 500 μl of nuclease-free water. The resuspended bacterial solution was applied onto the gelatine-coated mica sheet using a pipette tip. The sample was allowed to dry for 15 minutes before being subjected to AFM analysis. The AFM imaging was performed using a WITec alpha 300 R system. Images were acquired with different resolutions, including 1 μm, 2 μm, and 6 μm, to capture a range of details and surface features of interest.

## 3. Results

### 3.1. Enrichment and isolation of Methanotrophic strains

Methanotrophic bacteria were selectively enriched using NMS (Nitrate Mineral Salt) broth supplemented with methane as the sole carbon source, as described in the methodology section. The soil samples were first dissolved in distilled water and subsequently filtered. Serial dilutions were then performed using fresh NMS broth medium until a dilution factor of 10^−4^ was reached.

These serially diluted samples were incubated at 30°C and 150 RPM for 4 to 5 days, allowing sufficient time for proper bacterial growth. Following the incubation, an inoculum of 100 μl from the 10^−4^ diluted culture was spread onto NMS agar plates, resulting in the formation of isolated colonies. From these isolated colonies, further streaking was performed on additional NMS agar plates. Genomic DNA isolation was carried out from these streaked colonies, and subsequent PCR amplification was performed to identify the specific types of methanotrophs present. In the case of the samples obtained from the Oil & Natural Gas Field (ONGF), the isolated strain was identified as belonging to the *Methylocaldum* genus. Conversely, the strain isolated from the paddy field (PF) belonged to the *Methylomonas* genus.

### 3.2. Molecular identification of selected isolates

In methanotrophic bacteria, the 16S rRNA gene is approximately 1500 base pairs in length, while the pMMO (methanotrophic-specific gene) is around 900 base pairs long (22). Accordingly, the streaked isolated colonies obtained from ONGF site-2 at a dilution of 10^−4^ (**Supplementary Fig. 1b**) were used to inoculate NMS broth cultures for the isolation of genomic DNA. However, only the colonies marked with a red asterisk displayed proper growth. Genomic DNA was isolated from these specifically marked colonies **(Supplementary Fig. 1b)**, followed by amplification of the 16S rRNA gene and pMMO gene, as depicted in **Fig. 2a**. All the marked colonies selected from ONGF site-2 at a dilution of 10^−4^ demonstrated the presence of both the 16S rRNA gene and the pMMO gene, indicating the presence of methanotrophic bacterial species.

**Fig 1:**
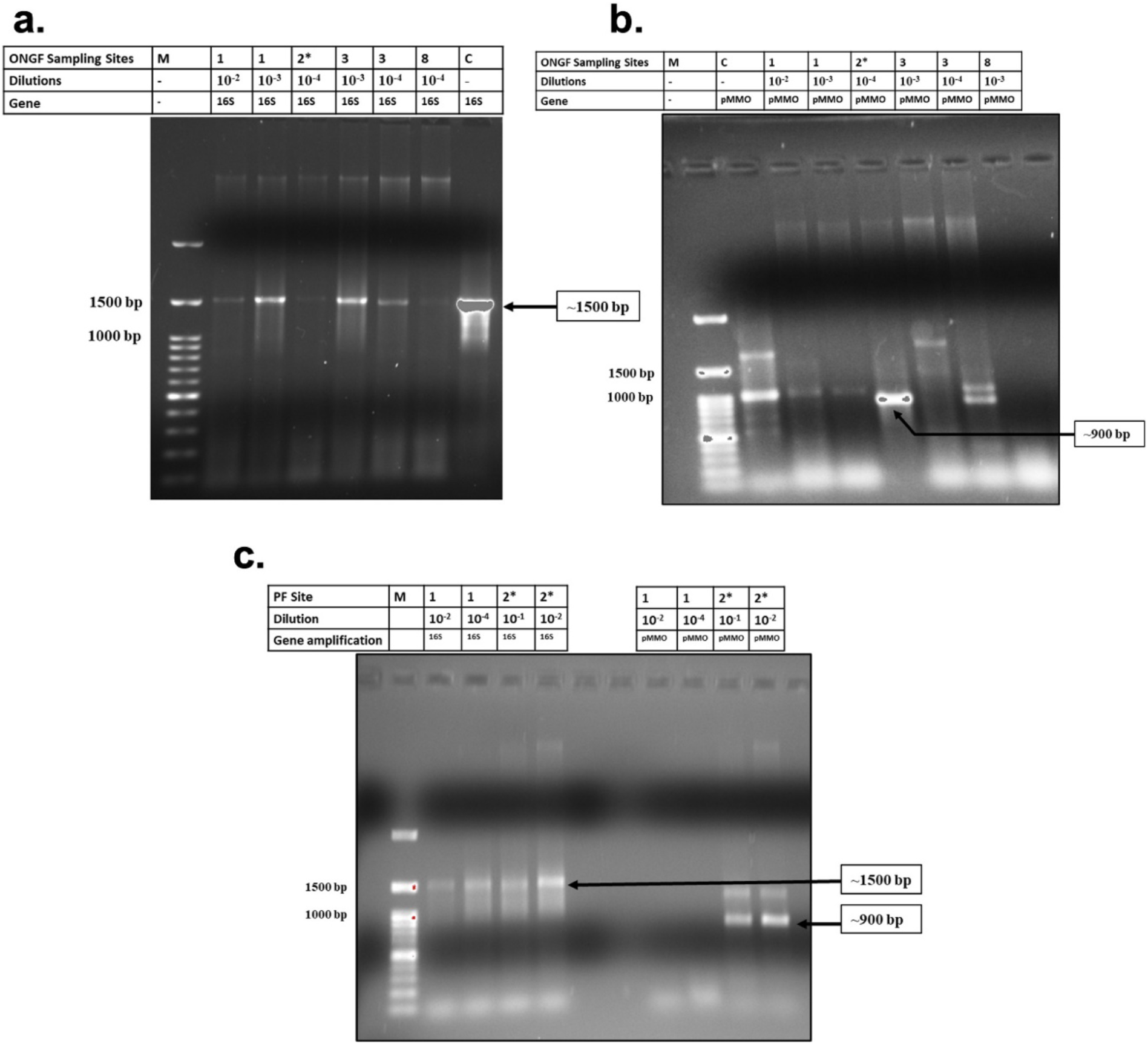
**(a)** PCR amplified products of genomic DNA isolated from different dilutions of ONGF sites using 16S rRNA gene-specific primers. M indicates the 100 base pairs DNA ladder, and C is the control (*Methylocystis parvus)*. **(b)** PCR amplified products of genomic DNA isolated from different dilutions of ONGF sites using pMMO gene-specific primers. M indicates the 100 base pairs DNA ladder, and C is the control (*Methylocystis parvus)*. **(c)** PCR amplified products of genomic DNA isolated from different dilutions of PF sites using 16S rRNA and pMMO gene-specific primers. M indicates the 100 base pairs DNA ladder.

**Fig 2:**
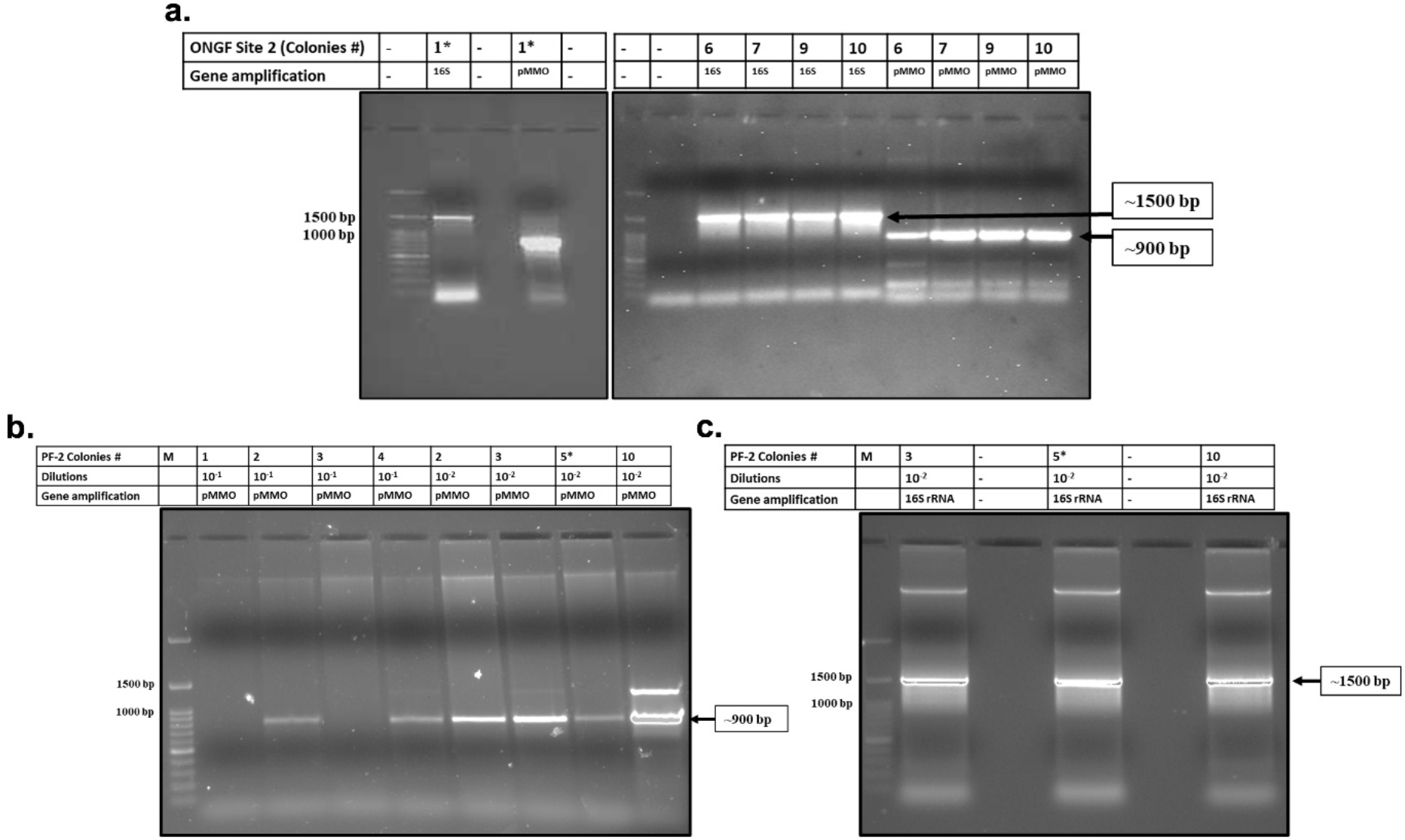
**(a)** Amplification of 16S rRNA and pMMO gene using respective primers for the genomic template DNA obtained from the isolated colonies of ONGF site-2 (dilution 10^−4^). **(b)** Amplification of pMMO gene using specific pMMO primers for the genomic template DNA obtained from the isolated colonies of PF site-2 (dilution 10^−1^ and 10^−2^). **(c)** Amplification of 16S rRNA gene using respective primers for the genomic template DNA obtained from the isolated colonies of ONGF site-2 (dilution 10^−2^). *Samples selected for further phylogenetic and morphological studies.

Likewise, the streaked isolated colonies marked with a red asterisk from PF site-2 at a dilution of 10^−2^ (**Supplementary Fig. 1d**) were examined for the amplification of the 16S rRNA gene (**Fig. 2b**), and pMMO gene (**Fig. 2c**). All the marked colonies selected from PF site-2 at a dilution of 10^−2^ exhibited the presence of both the 16S rRNA gene and the pMMO gene, suggesting the presence of methanotrophic bacterial species.

### 3.3. Nucleotide sequence accession number

The 16S rRNA gene sequence of four ONGF-isolated methanotrophic sp. (Isolate 1, 6, 9, and 10) and one PF-isolated methanotrophic sp. (Isolate 5) have been deposited to the GenBank database. Among all four isolates of ONGF, only Isolate-1 was further studied for their phylogenetic relationship and morphology, as all four isolates showed nucleotide similarity to the same two reference sequences with accession numbers KY621548.1 and EU275146.1. Detailed information about the submitted sequence has been provided in **Table 3**.

**Table 3:** Nucleotide sequence submitted in GenBank database.

| Submitted 16S rRNA nucleotide sequence of bacterial sp. isolated in this study |  |  |  | Homology to other bacterial species (%), Accession no. |
| --- | --- | --- | --- | --- |
| Sample | Strain | Submission Id | Accession number |  |
| ONGF Isolate 1 | SSM3AK117ISOLATE1_16S | SUB12362364 | OP955938 | - *Methylocaldum gracile* strain VKM 14L 16S rRNA gene, partial sequence (97.25%), **NR_026063.1**<br>- *Methylocaldum marinum* DNA, complete genome, strain S8 (96.12%), **AP017928.1** |
| ONGF Isolate 6 | SSM3AK117ISOLATE6_16S | SUB12368462 | OP954828 | - *Methylocaldum gracile* strain VKM 14L 16S rRNA gene, partial sequence (98.54%), **NR_026063.1**<br>- *Methylocaldum marinum* DNA, complete genome, strain S8 (97.30%), **AP017928.1** |
| ONGF Isolate 9 | SSM3AK117ISOLATE9_16S | SUB12368465 | OP954854 | - *Methylocaldum gracile* strain VKM 14L 16S rRNA gene, partial sequence (97.42%), **NR_026063.1**<br>- *Methylocaldum marinum* DNA, complete genome, strain S8 (96.31%), **AP017928.1** |
| ONGF Isolate 10 | SSM3AK117ISOLATE10_16S | SUB12368470 | OP954856 | - *Methylocaldum gracile* strain VKM 14L 16S rRNA gene, partial sequence (98.39%), **NR_026063.1**<br>- *Methylocaldum marinum* DNA, complete genome, strain S8 (97.37%), **AP017928.1** |
| PF Isolate 5 | SSM3AK117_PCPF_PII_10-2_5_16 | SUB12367512 | OP957340 | - *Methylomonas koyamae* strain Fw12E-Y DNA, complete genome (84.99%), **AP019777.1**<br>- *Methylomonas koyamae* strain Fw12E- Y, 16S rRNA gene, partial sequence (84.99%), **NR_113033.1** |

### 3.4. Phylogenetic analysis

To determine the impact of using 16S rRNA and pmoA genes as main molecular markers for defining new methanotrophic taxa, phylogenetic analyses were performed using sequences from 14 Query sequences available in the NCBI database for the generation of phylogenetic trees using MEGA-X. The trees were built up using the MUSCLE algorithm with 1000 bootstrap replicates.

In the case of the ONGF isolated strain, the 16S rRNA-based phylogenetic tree showed that the *Methylocaldum* and *Methylococcus* are very much related to the query sequence (**Fig. 3a**). However, pmoA-based phylogenetic tree showed that the query sequence was 86% similar to the *Methylocaldum* group of bacterial species. Despite this, *Methylomarinovum caldicuralii, Methylohalobius crimeensis, Methylomarinum vadi, Methylovulum miyakonense, Methylotuvimicrobium japanese, Methylococcus capsulatus, and Methylococcus geothermmalis* also showed similarity to the query sequence (**Fig. 3b**). By analysing both 16S rRNA and pMMO sequence-based phylogenetic trees, the isolated strain was considered as putative *Methylocaldum* species.

**Fig 3:**
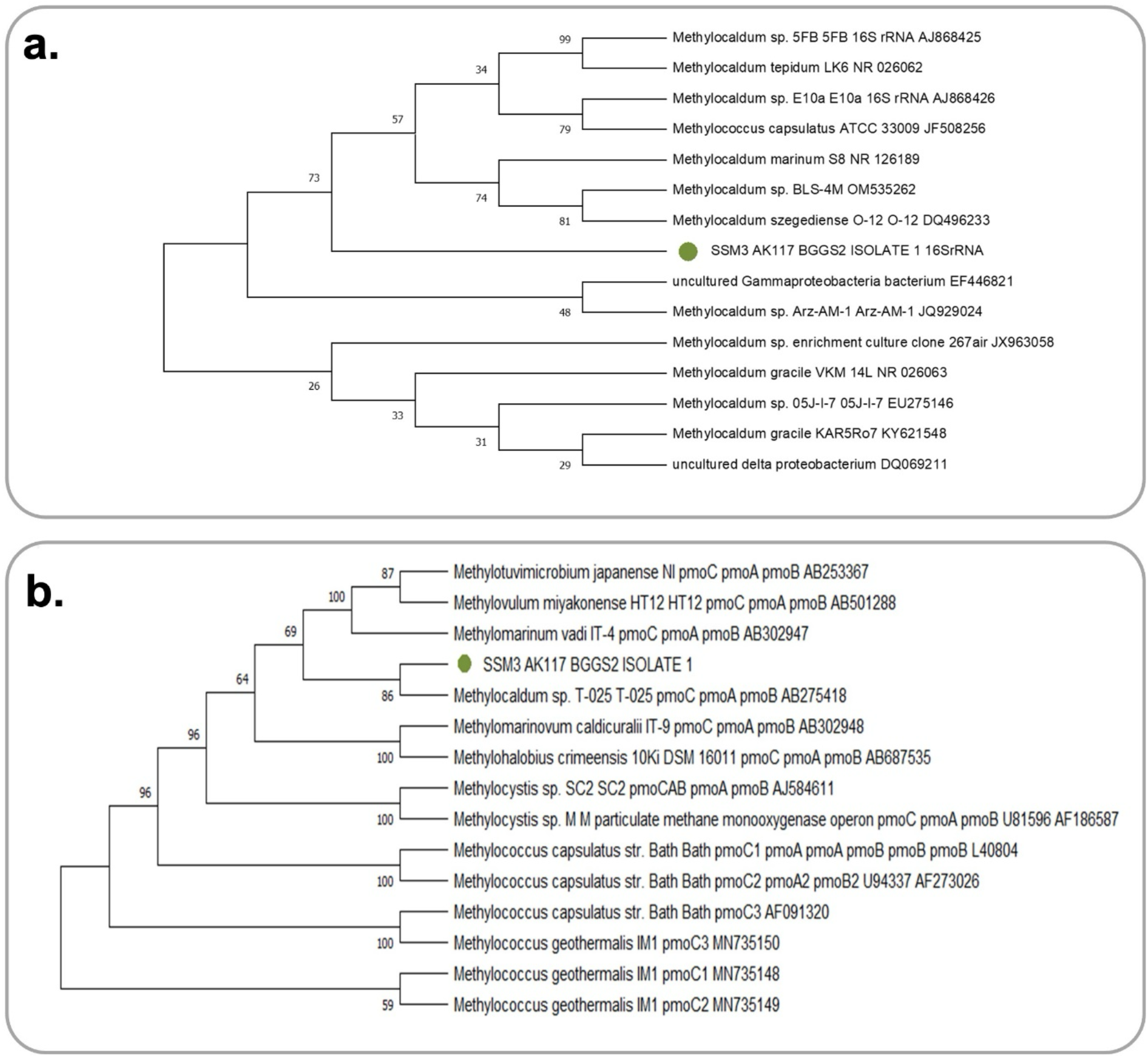
The phylogenetic relationship of 14 partial methanotrophs sequences compared with the isolated ONGF soil sample **(a)** Borholla GGS 2_16S rRNA sequences and **(b)** Borholla GGS 2_pMMO sequences (green marked). This phylogenetic tree is generated by the Maximum Likelihood method using the MUSCLE algorithm with 1000 bootstrap replicates using the MEGA X version 11 building program.

In the case of the PF isolated strain, the 16S rRNA gene sequence amplified from this isolated strain showed 78% similarity to uncultured *Methylomonas* (**Fig 4a**). The pMMO gene sequence amplified from this strain showed 97% similarity with the *Methylomonas* sp. However, it was also phylogenetically related to *Methylomonas methanica* and *Methylomarinum vadi* (**Fig 4b**). By analysing 16S rRNA and pMMO sequence-based phylogenetic trees, the isolated strain was considered as a putative *Methylomonas* species.

**Fig 4:**
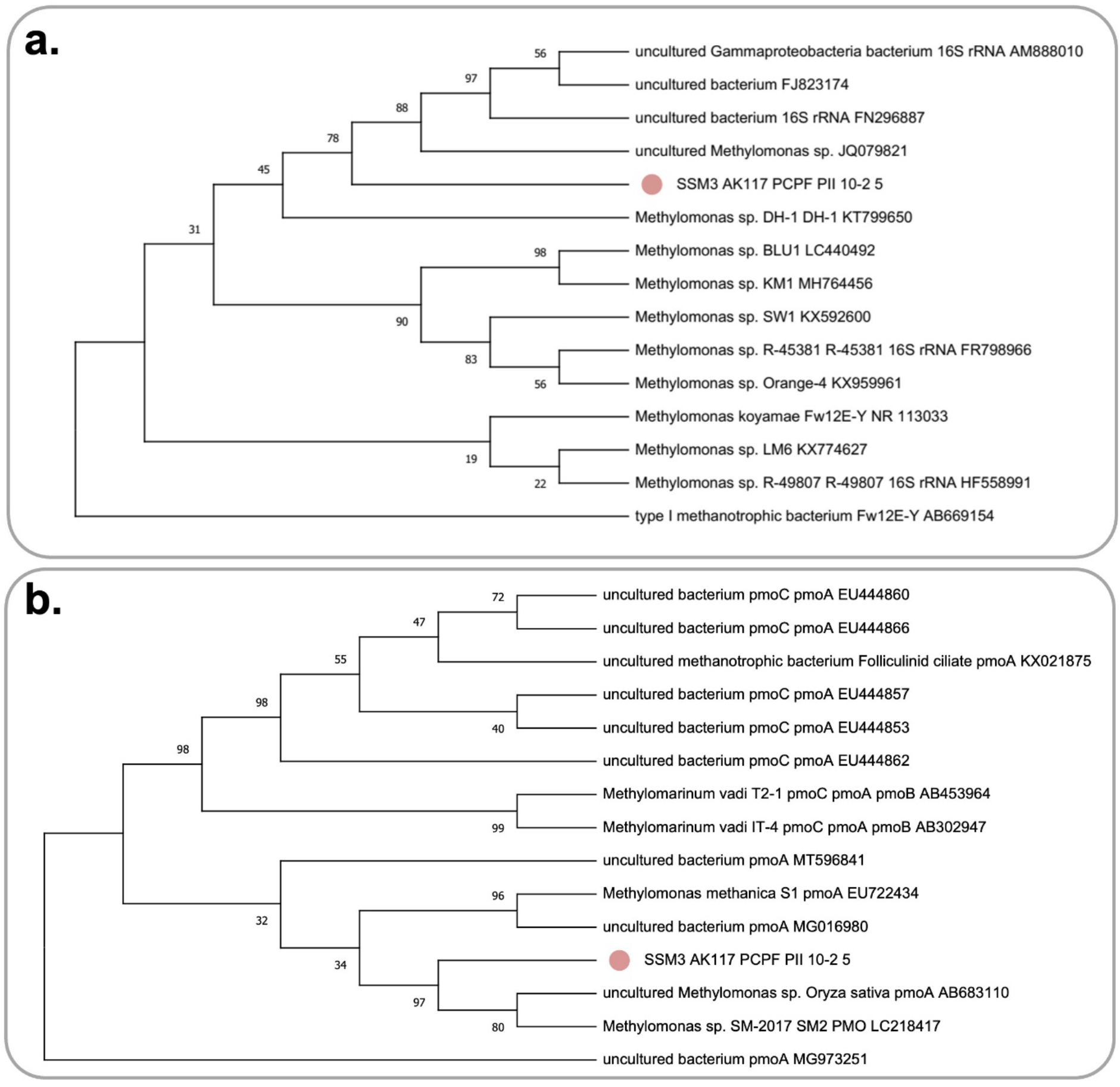
The phylogenetic relationship of 14 partial methanotrophs sequences compared with our purely isolated oil field soil sample **(a)** PCPF_16S rRNA sequences and **(b)** PCPF_pMMO sequences (pink marked). This phylogenetic tree is generated by the Maximum Likelihood method using the MUSCLE algorithm with 1000 bootstrap replicates using the MEGA X version 11 building program.

### 3.5. Morphological characterisation

The bacterial species isolated from the ONGF site-2 showed similarity to the *Methylocaldum* sp., which tends to have either rod pleomorphic or cocci shape (23). Using AFM, the shape of isolated species isolated from ONGF site-2 was also observed as a cocci shape (**Fig. 5a-b, Supplementary Fig 2 a-b**), confirming its belonging to the *Methylocaldum* species. The bacterial species isolated from the PF site-2 showed similarity to the *Methylomonas* sp., which tends to have a rod shape (24). Using AFM, the shape of isolated species isolated from PF site-2 was also observed as rod shape (**Fig. 5c-d, Supplementary Fig 2 c-d**), confirming its belonging to the *Methylomonas* species.

**Fig 5:**
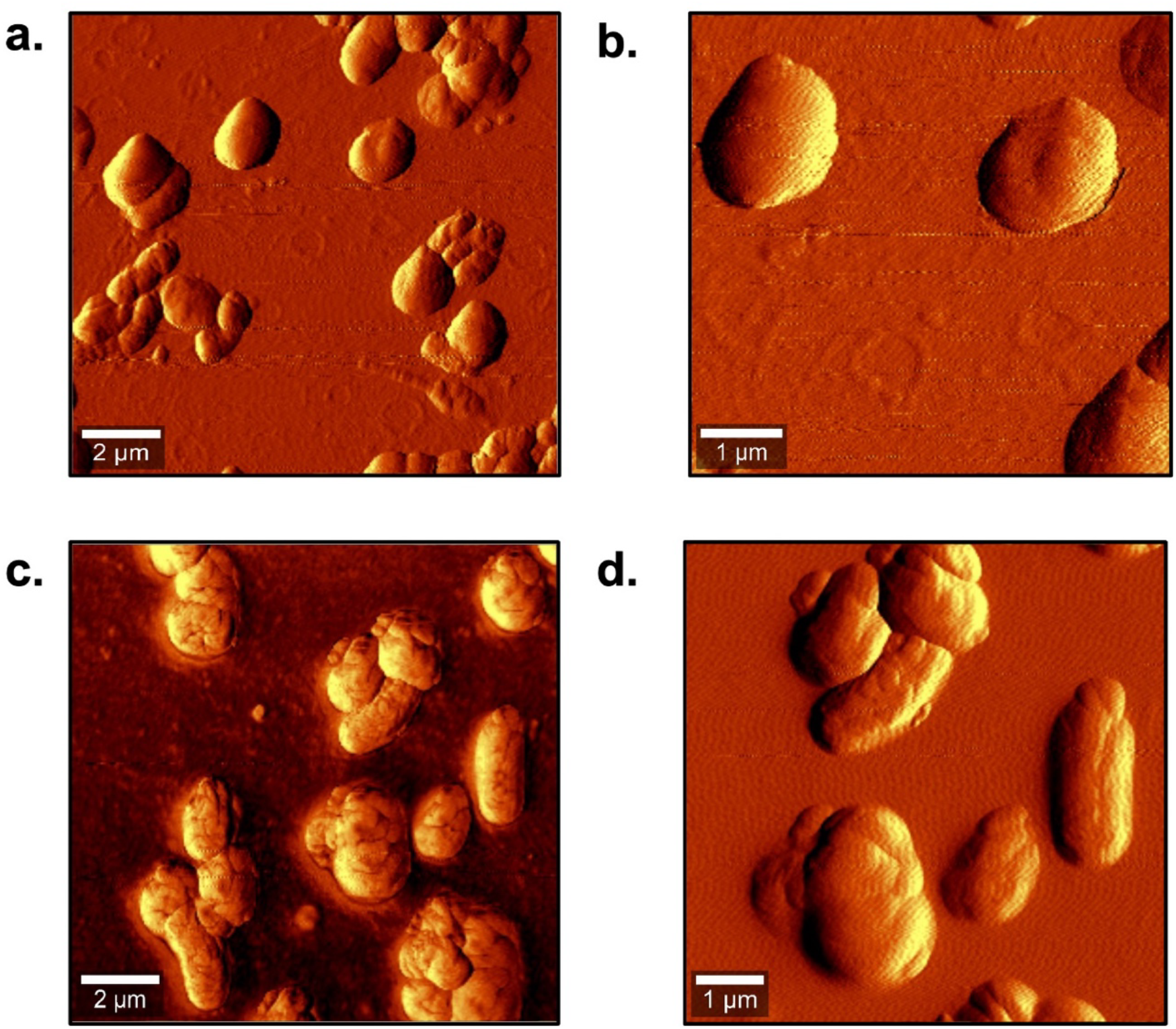
Atomic Force Microscopy (AFM) images of isolated bacterial species. **(a-b)** *Methylocaldum* sp. with cocci shape isolated from ONGF site 2. **(c-d)** *Methylomonas* sp. with rod shape isolated from PF site-2.

## 4. Discussion

To enrich and isolate methanotrophs, specific culture media such as NMS supplemented with methane gas as the sole carbon source are required. In this study, we aimed to identify and characterise methanotrophic bacteria from two geographically isolated sites. The soil samples were serially diluted and subjected to 16S rRNA and pMMO gene amplification for identification. Additionally, the isolated methanotrophs were examined for their morphological characteristics.

Based on the analysis of 16S rRNA and pMMO gene sequences, we observed that the four isolates from ONGF site-2 and one isolate from PF site-2 were closely related to various cultured species of methanotrophic bacteria. Previous studies utilizing molecular detection methods have also revealed the presence of these methanotrophic species in the soil. For the ONGF isolates, the top 100 selected species included 16 known methanotrophic species, exhibiting sequence homology ranging from 91.82% to 97.25%, as identified in the NCBI database. A phylogenetic tree depicting the relationship between the four ONGF isolates and other reference sequences in the NCBI database was constructed using the MOLE-BLAST tool (available online at https://blast.ncbi.nlm.nih.gov/moleblast/moleblast.cgi). Based on their lower similarity (91.82% to 97.25%) to closely related relatives, all four ONGF isolates were considered to belong to the *Methylocaldum* genus (**Supplementary Fig 3**). Regarding PF isolate 5, among the top 100 selected species, 40 were known methanotrophic species, showing sequence homology between 79.72% and 84.99%. This isolate was classified as a member of the *Methylomonas* genus, given its low similarity (79.72% to 84.99%) to closely related relatives (**Fig. 4**). The identification of isolated strains from ONGF, including Isolate 1 (SSM3AK117ISOLATE1_16S) (**Supplementary Fig. 4**), Isolate 6 (SSM3AK117ISOLATE6_16S) (**Supplementary Fig. 5**), Isolate 9 (SSM3AK117ISOLATE9_16S) (**Supplementary Fig. 6**), and Isolate 10 (SSM3AK117ISOLATE10_16S) (**Supplementary Fig. 7**), was further validated by applying the “Limit to sequence from type material” filter during BLAST-NCBI homology searching, confirming their homology to *Methylocaldum* species. Similarly, the same filter was applied for PF-isolated strains, specifically Isolate 5 (SSM3AK117_PCPF_PII_10-2_5_16) (**Supplementary Fig. 8**), which showed homology to *Methylomonas* species.

To reinforce the molecular detection and phylogenetic observations, one of the ONGF isolates (Isolate 1) and the only available PF isolate (Isolate-5) were selected for morphological studies. The initial phylogenetic analysis results supported the identification of *Methylocaldum* species from ONGF site-2 and *Methylomonas* species from PF site-2. These findings were further corroborated by microscopic examination. Previous reports have described the cellular shapes of *Methylocaldum* species as either rod-shaped, pleomorphic, or cocci (Takeuchi et al., 2014b), and the ONGF isolates exhibited a cocci shape as well using AFM. *Methylomonas* species, on the other hand, typically display a rod-shaped morphology (24), which was also observed in the PF isolates. Therefore, these observations support that the bacterial species isolated from the ONGF site-2 and PF site-2 belong to the *Methylocaldum* and *Methylomonas* genera, respectively.

## 5. Conclusion

Methanotrophs growing in the methane emission sites such as oil-natural gas fields and paddy fields used in this study are majorly involved in methane to methanol conversion. These microorganisms play a crucial role in methane reduction naturally and may also be helpful for the industrial production of biodiesel. In this study, two methanotrophic species belonging to the *Methylocaldum* and *Methylomonas* were identified from the oil-natural gas field and paddy field, respectively, using PCR amplification. The length of the 16S rRNA gene and pMMO gene amplified from both the isolated species were ~1500 bps and ~900 bps, indicating the presence of methanotrophic bacteria. Further microscopic studies of ONGF Isolated and PF Isolated species also revealed their resemblance to bacterial species from *Methylocaldum* and *Methylomonas* genera.

## Supporting information

https://docs.google.com/document/d/14HtZABkpuHmy2GxM6QN563cRdqHJOkMm/edit?usp=drive_link&ouid=100938542357473714946&rtpof=true&sd=true

## Acknowledgements

The authors thank Prof. V.S Moholkar, Dr Lepakshi Borbora and Ms Aradhana Priyadarsini, INBIGS & ONGC Jorhat, Assam, for helping and providing facilities in collecting the soil samples from the Jorhat, Assam sites. Funding support was provided by DBT Project BT/PR25338/NER/95/1147 /2017, dated 29/09/2018. The authors also want to acknowledge AIRF-JNU, New Delhi, for providing an atomic force microscopy (AFM) facility. Author Akanksha Verma wants to acknowledge DST-INSPIRE for providing financial support during five years of PhD tenure.

## Declarations

### Author Contributions

AV is involved in Conceptualization, Methodology, Data Curation and analysis, Writing-original draft, and Editing; SSM is involved in the Supervision, Conceptualization, Validation, Writing-review & editing. All authors made important contributions to the manuscript and approved publication.

