## Supplementary material for "Differences in Methanotrophic Community Structure in Two Methane-Rich Habitats: Oil Natural Gas Field & Paddy Field": https://docs.google.com/document/d/14HtZABkpuHmy2GxM6QN563cRdqHJOkMm/edit?usp=drive_link&ouid=100938542357473714946&rtpof=true&sd=true

**Supplementary File 1**

**Differences in Methanotrophic Community Structure in Two Methane-Rich Habitats: Oil Natural Gas Field & Paddy Field**

Akanksha Verma^1^, S.S. Maitra^1^

*^1^School of Biotechnology, Jawaharlal Nehru University, New Delhi-110067, India*

---------------------------------

**Akanksha Verma**

ORCID ID: 0000000328664012

***Corresponding author:** S. S. Maitra

ORCID ID: 0000000239557004

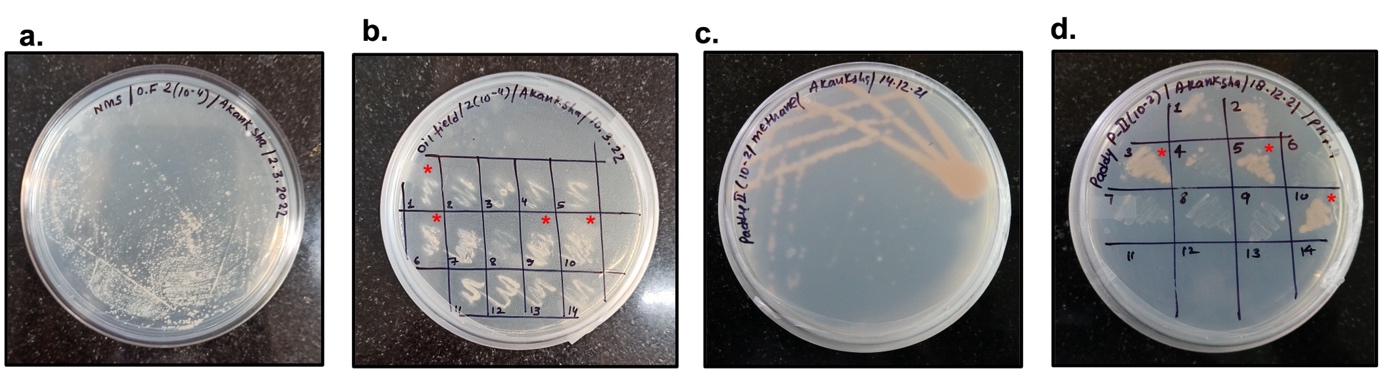

**Supplementary Fig 1:** **(a)** Bacterial colonies were obtained in the presence of methane on the NMS agar plate from the culture of ONGF site-2 (dilution 10^-4^). **(b)** Isolated colonies from the culture of ONGF site-2 (dilution 10^-4^) streaked on the NMS agar plate. Colonies marked with a red * sign were further selected for the identification of methanotrophic strains using 16S rRNA and pMMO gene sequencing. **(c)** Quadrant streaking of bacterial culture from PF site-2 (dilution 10^-2^) on NMS agar plate and grown in the presence of methane to obtain isolated colonies. **(d)** Isolated colonies from the culture of PF site-2 (dilution 10^-2^) streaked on the NMS agar plate. Colonies marked with a red * sign were further selected for the identification of methanotrophic strains using 16S rRNA and pMMO gene sequencing.

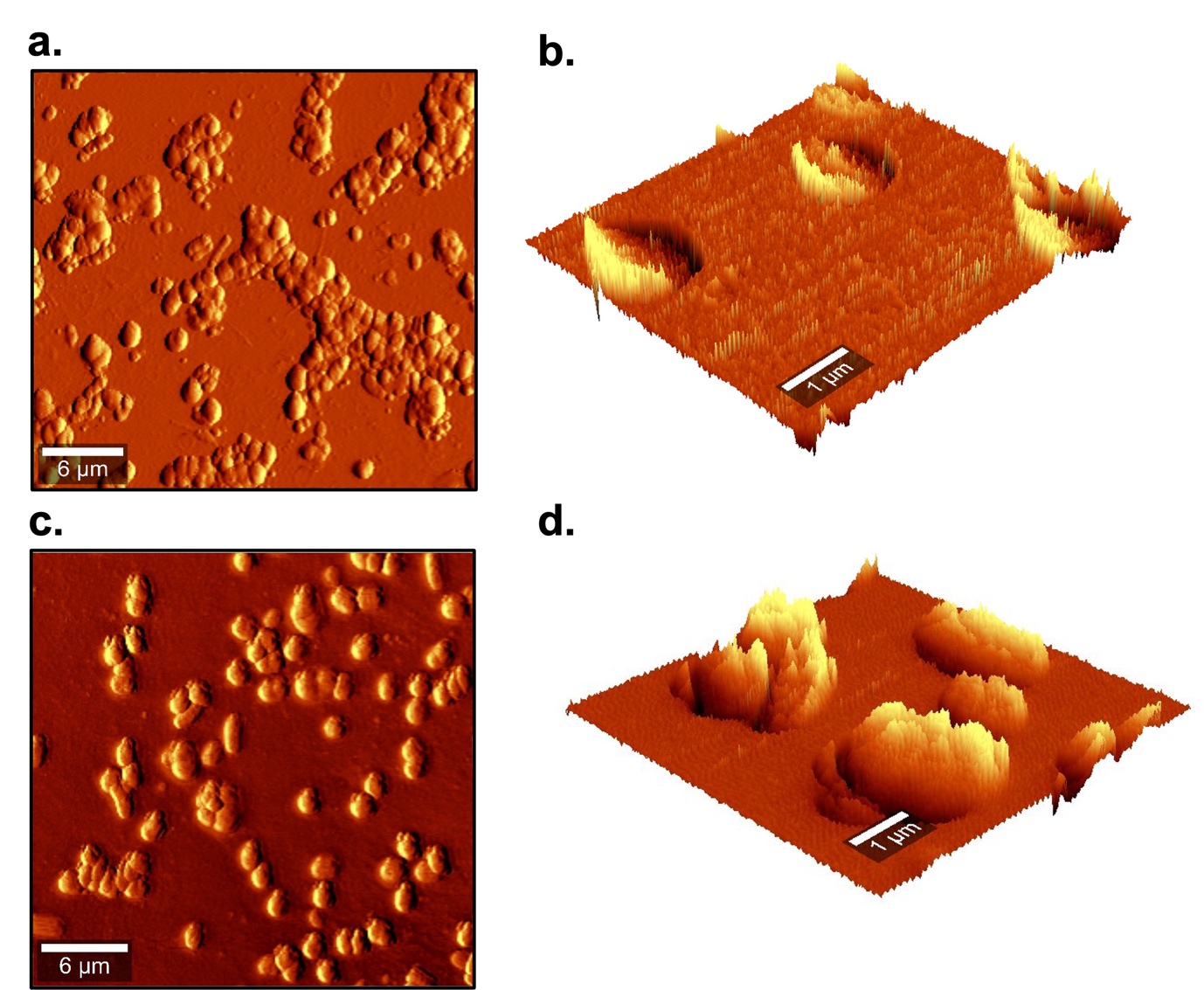

**Supplementary Fig 2**: Atomic Force Microscopy (AFM) images of isolated bacterial species. **(a-b)** *Methylocaldum* sp. with cocci shape isolated from ONGF site 2. **(c-d)** *Methylomonas* sp. with rod shape isolated from PF site- 2.

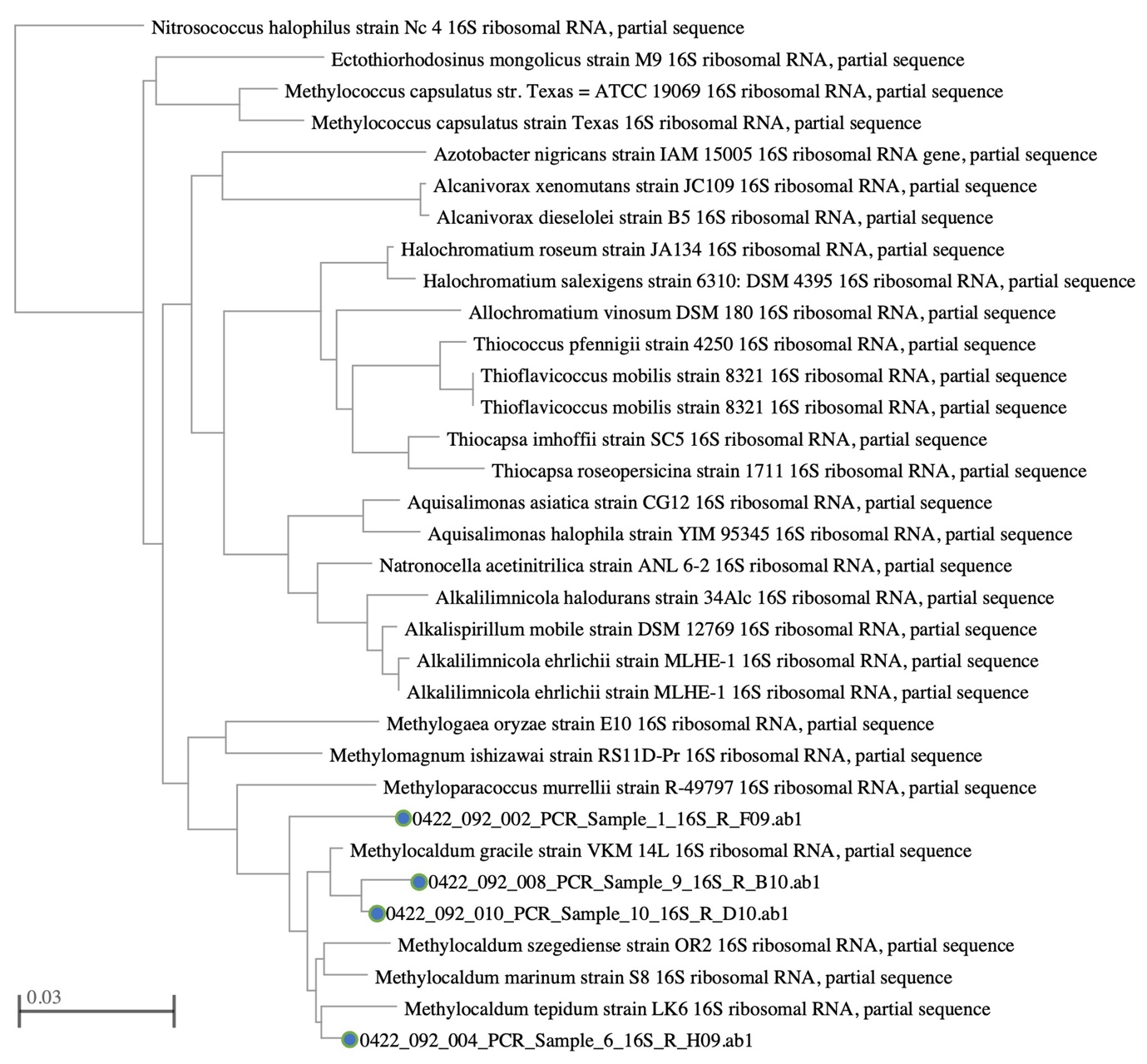

**Supplementary** **Fig 3**: The phylogenetic relationship of all four ONGF isolated species, showing similarity to the *Methylomonas* genus. The tree was constructed using the MOLE-BLAST online server.

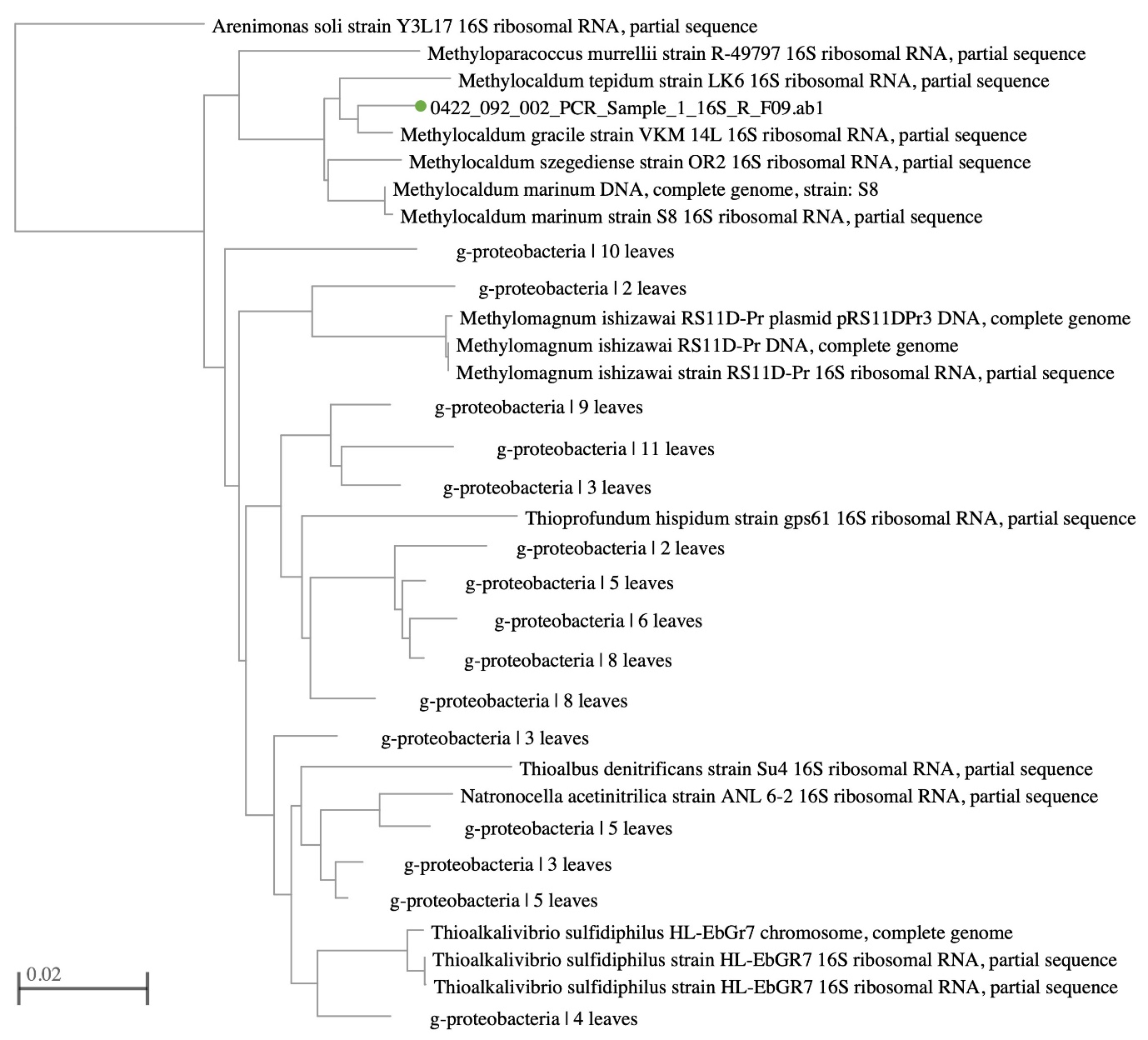

**Supplementary** **Fig 4**: The Phylogenetic relationship of ONGF Isolate 1 with species of higher similarity (In BLAST- NCBI ‘Limit to sequences from type material’ is filtered).

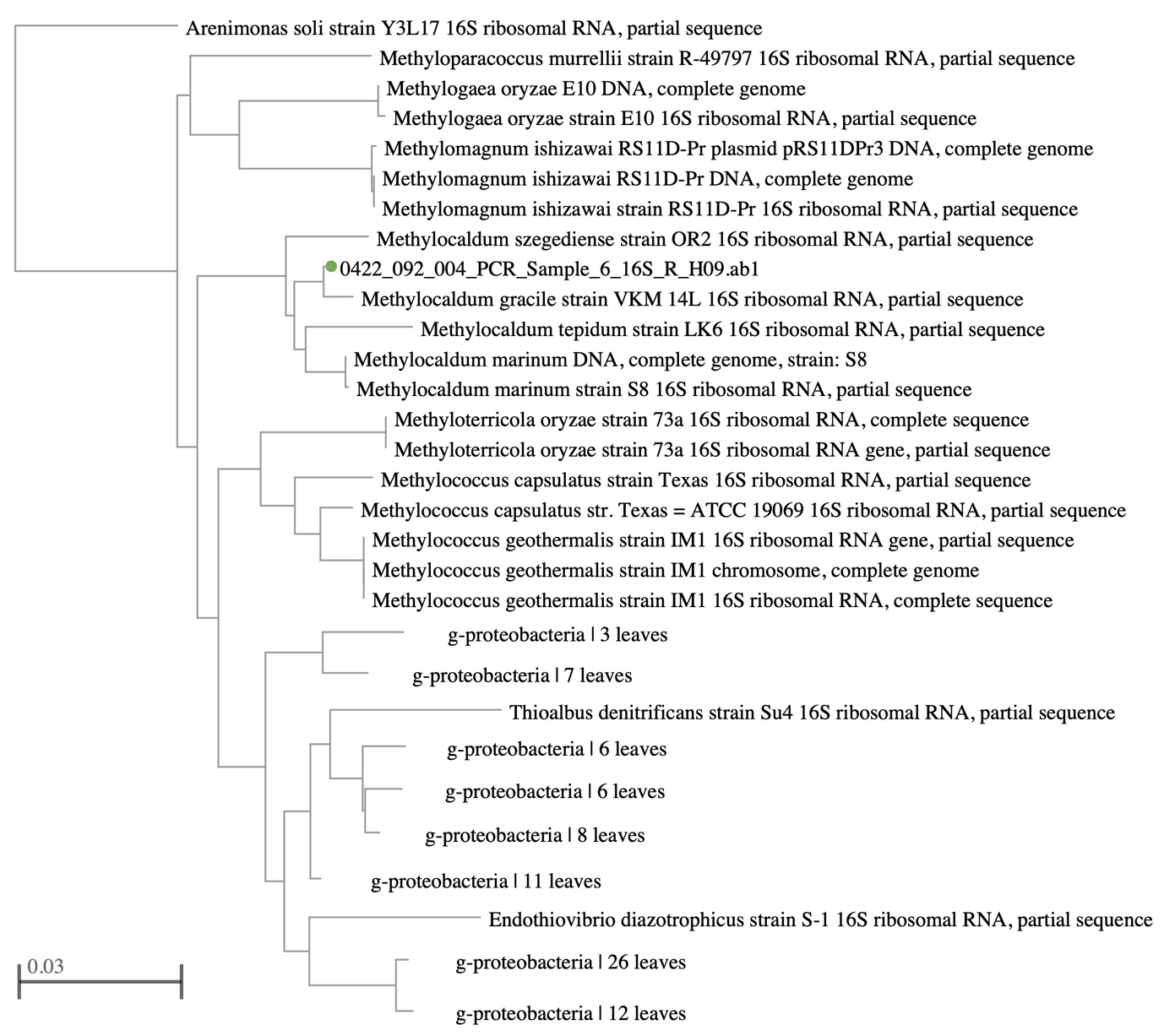

**Supplementary** **Fig 5**: The Phylogenetic relationship of ONGF Isolate 6 with species of higher similarity (In BLAST- NCBI ‘Limit to sequences from type material’ is filtered).

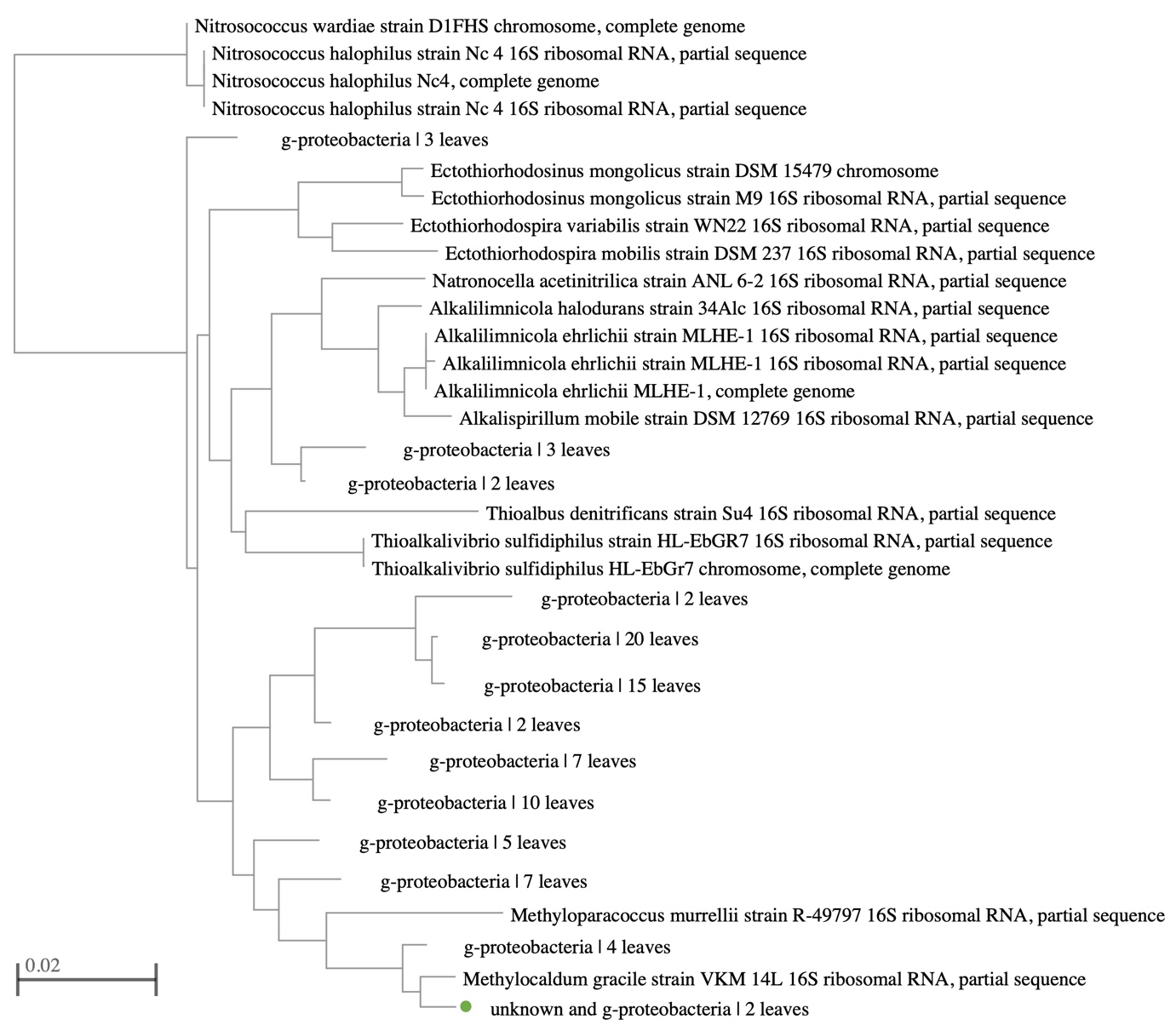

**Supplementary** **Fig 6**: The Phylogenetic relationship of ONGF Isolate 9 with species of higher similarity (In BLAST- NCBI ‘Limit to sequences from type material’ is filtered).

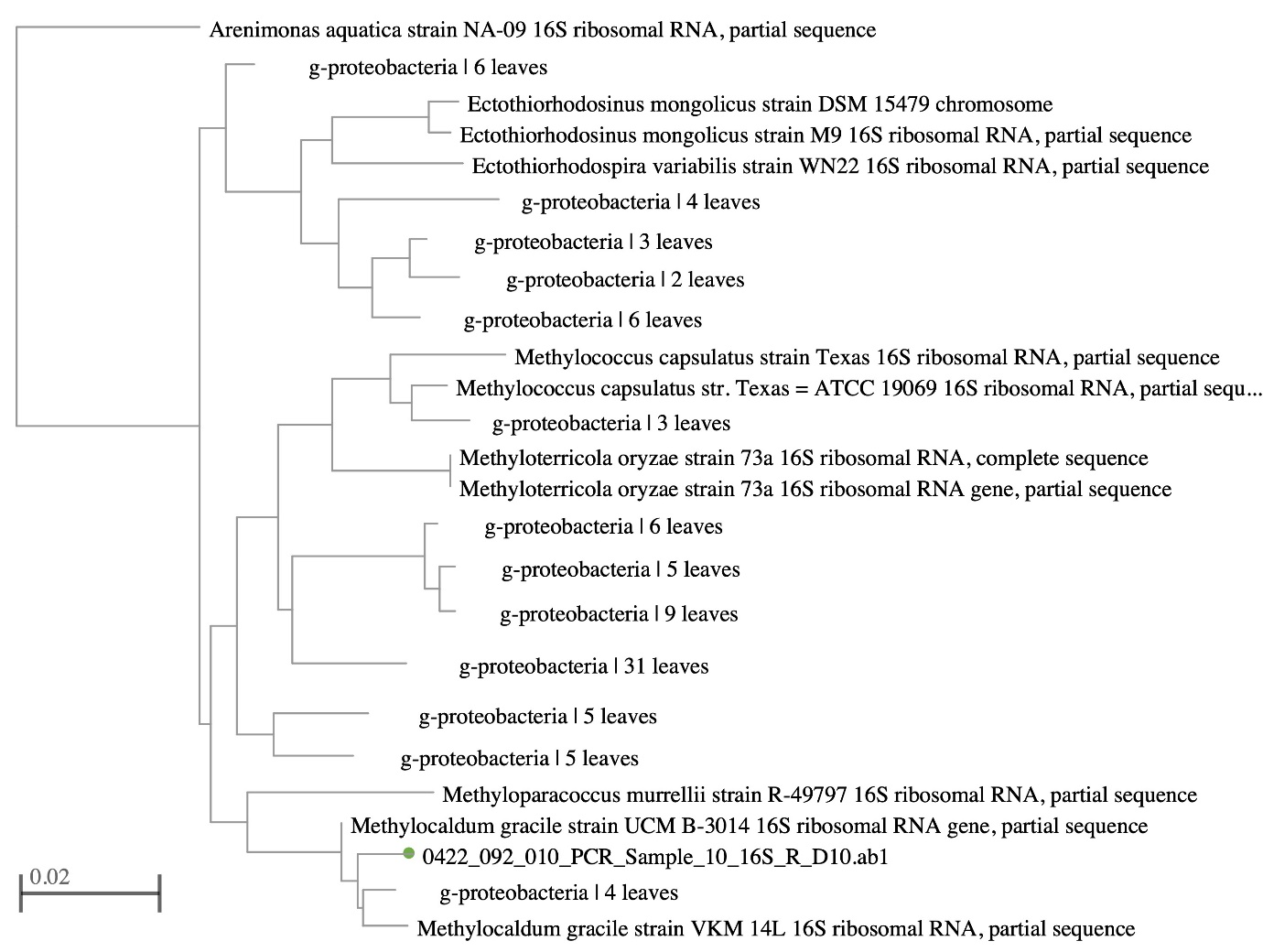

**Supplementary** **Fig 7:**  The Phylogenetic relationship of ONGF Isolate 10 with species of higher similarity (In BLAST- NCBI ‘Limit to sequences from type material’ is filtered).

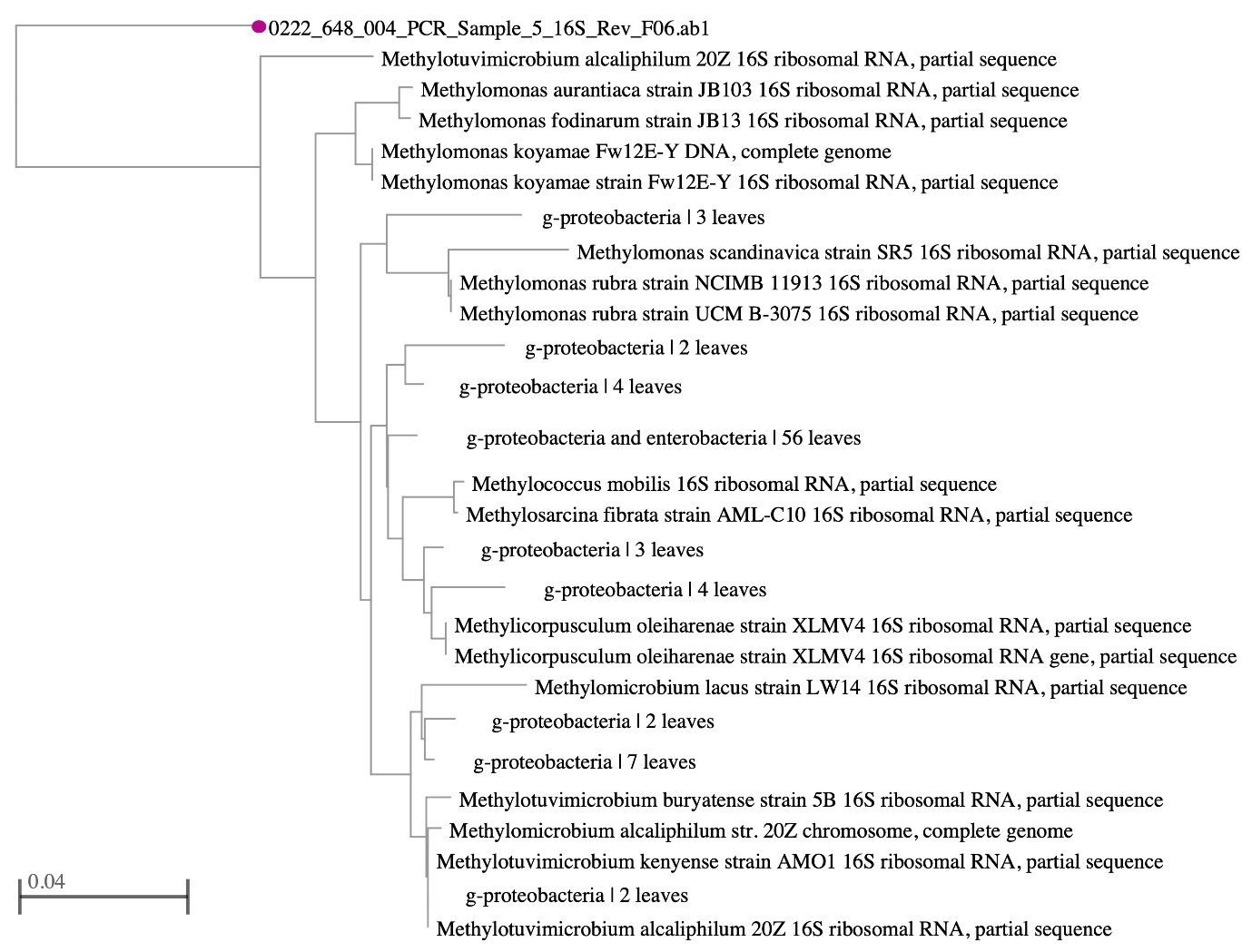

**Supplementary** **Fig 8**: The Phylogenetic relationship of PF Isolate 5 with species of higher similarity (In BLAST- NCBI ‘Limit to sequences from type material’ is filtered).
